# The developing Human Connectome Project fetal functional MRI release: Methods and data structures

**DOI:** 10.1101/2024.06.13.598863

**Authors:** Vyacheslav R. Karolis, Lucilio Cordero-Grande, Anthony N. Price, Emer Hughes, Sean P. Fitzgibbon, Vanessa Kyriakopoulou, Alena Uus, Nicholas Harper, Denis Prokopenko, Devi Bridglal, Jucha Willers Moore, Sian Wilson, Maximilian Pietsch, Daan Christiaens, Maria Deprez, Logan Z.J. Williams, Emma C. Robinson, Antonis Makropoulos, Seyedeh-Rezvan Farahibozorg, Jonathan O’Muircheartaigh, Mary A. Rutherford, Daniel Rueckert, A. David Edwards, Tomoki Arichi, Stephen M. Smith, Eugene Duff, Joseph V. Hajnal

## Abstract

Recent advances in fetal fMRI present a new opportunity for neuroscience to study functional human brain connectivity at the time of its emergence. Progress in the field however has been hampered by the lack of openly available datasets that can be exploited by researchers across disciplines to develop methods that would address the unique challenges associated with imaging and analysing functional brain in utero, such as unconstrained head motion, dynamically evolving geometric distortions, or inherently low signal-to-noise ratio. Here we describe the developing Human Connectome Project’s release of the largest open access fetal fMRI dataset to date, containing 275 scans from 255 fetuses and spanning the period of 20.86 to 38.29 post-menstrual weeks. We present a systematic approach to its pre-processing, implementing multi-band soft SENSE reconstruction, dynamic distortion corrections via phase unwrapping method, slice-to-volume reconstruction and a tailored temporal filtering model, with attention to the prominent sources of structured noise in the in utero fMRI. The dataset is accompanied with an advanced registration infrastructure, enabling group-level data fusion, and contains outputs from the main intermediate processing steps. This allows for various levels of data exploration by the imaging and neuroscientific community, starting from the development of robust pipelines for anatomical and temporal corrections to methods for elucidating the development of functional connectivity in utero. By providing a high-quality template for further method development and benchmarking, the release of the dataset will help to advance fetal fMRI to its deserved and timely place at the forefront of the efforts to build a life-long connectome of the human brain.

## 1. Introduction

Already at birth, brain activity appears to be organised into a “connectome” of distributed networks (Doria et al., 2010; Fitzgibbon et al., 2020; Gao et al., 2015) that underpin complex behaviours and cognitive functions later in life. Multiple lines of evidence now also point to the critical importance of the fetal period for healthy development (Bergman et al., 2007; Boukhris et al., 2016; Brown et al., 1995; Laplante et al., 2008; O’Donnell et al., 2009; Rakers et al., 2020; Sandman et al., 2012; Skranes & Løhaugen, 2022; Zerbo et al., 2015). There is thus an increasing need for data on how functional connections become established in the prenatal brain.

Advances in fetal fMRI present an opportunity for neuroscientific studies of the functional human brain at the time of its emergence (Ferrazzi et al., 2014; Schöpf et al., 2012). Imaging an in-utero brain, however, poses numerous unique challenges (Ferrazzi et al., 2014; Seshamani et al., 2016; Sobotka et al., 2022; Taymourtash et al., 2019). Unconstrained motion, non-rigid maternal tissues surrounding the fetal brain, a greater distance between the head and receiver coil are among the MRI-adverse factors that lower signal-to-noise ratio (SNR) and increase the level of structured artifacts. Motion-free periods are empirically rare in *in utero* fMRI acquisitions given that motion associated with maternal respiration and fetal movement itself systematically induce fetal head displacements. When extreme, these displacements may prohibit the reconstruction of particular volumes in the timeseries (Sobotka et al., 2022; Taymourtash et al., 2021). When mild to moderate, they can still cause significant spin-history effects (Ferrazzi et al., 2014; Seshamani et al., 2016), manifested by spatially non-stationary image intensity changes (*travelling waves*), not readily amenable to established methods of data temporal filtering (Griffanti et al., 2014; Salimi-Khorshidi et al., 2014). The interaction between the magnetic properties of the moving fetal head and the adjacent maternal tissues induces dynamic changes in the “static” magnetic field (B0) inhomogeneity that results in temporally evolving distortion of fetal brain geometry (Cordero-Grande et al., 2018a). The high contrast of maternal tissues may also induce leakage artifacts in the multi-band (MB) sensitivity encoded (SENSE) MRI data reconstruction. Overall, the *in utero* setting introduces multiple challenges to be navigated *en route* to artifact-controlled characterisation of emerging brain functional connectivity.

Finding solutions to these challenges, as well as the development of tailored methods for fetal brain connectivity analyses, requires a community-wide effort. Progress in this direction however is being hampered by the lack of openly available datasets that could be exploited by researchers across disciplines. The developing Human Connectome Project (dHCP) closes this gap by releasing the first open-access and largest-to-date fetal fMRI dataset (Data and code availability, Resource 1) of 275 scans from 255 individuals (gestational age (GA): 20.86 - 38.29 weeks), processed using tailored methods and accompanied by an advanced registration infrastructure between imaging and template spaces. The dataset complements the open-access dataset of neonatal fMRI data (Fitzgibbon et al., 2020), together allowing for detailed investigations of connectivity in the perinatal brain. In this paper we set forth details of this endeavour to process the fetal fMRI data from the stage of frequency-to-image reconstruction all the way to the level when they can be utilised for group-level analyses. To aid future method benchmarking, we make available the outputs from all the main pre-processing stages. This allows for various levels of data exploration by the imaging and neuroscientific community, starting from the development of robust pipelines for anatomical, distortion and temporal corrections to the analyses investigating the development of the prenatal functional connectivity. The aim of this paper is threefold: 1) describe the organisation of the dHCP fetal fMRI release data, 2) provide detailed descriptions and motivations for the pre-processing methods implemented in the released data; 3) demonstrate its capacity for performing group-level analyses. We interleave descriptions of methods and results for a clearer exposition of analytical approaches and their contribution to the improvement of data quality.

## 2. Methods and Results

### 2.1. Acquisition parameters and data sample

Participants were prospectively recruited as part of the dHCP, a cross-sectional open science initiative funded by the European Research Council. Resting-state fMRI data were acquired with a Philips Achieva 3T system (Best, NL) using a 32-channel cardiac coil at St Thomas’ Hospital London. Scanner software was R3.2.2 with a custom patch. Each data set consists of 350 volumes (48 slices each), acquired using a single-shot echo planar imaging (EPI) (TR/TE = 2200/60) sequence, with slice matrix = 144 x 144, isotropic resolution = 2.2 mm, MB factor = 3, and SENSE factor = 1.4 (Price, 2019). Brains of all fetuses were reported by a neuroradiologist as showing appropriate appearances on the T2-weighted anatomical scan for their GA with no acquired lesions or congenital malformations of clinical significance.

A total of 277 completed fetal fMRI scans were acquired. Two were excluded from public data release due to poor data quality across all modalities (T2-weighted, diffusion, and fMRI). The remaining 275 fMRI sessions were acquired from 255 unique individual subjects (137 male, 116 female, 2 unrecorded, GA: 20.86 - 38.29 weeks). The mothers of 77.25% babies identified themselves as white, 2.75% as black, 13.33% of any Asian origin (including South East Asia), 3.53% of any mixed origin, 2.75% of other unspecified origin, and 1 case refusing to provide this information. The details of ethnicity as well as mother’s medical, obstetric and mental health information are made available with the release (Data and code availability, Resource 1). For comparison, the average numbers for Greater London and South East England, i.e., recruitment area for the study, are: white - 70.1%, black - 7.95%, Asian - 13.85%, mixed - 4.25%, other - 3.9% (https://www.ethnicity-facts-figures.service.gov.uk/uk-population-by-ethnicity/national-and-regional-populations/regional-ethnic-diversity/latest/).

Following an initial visual assessment, 1 scan was excluded from further processing due to an incomplete field-of-view. With one exception, all fMRI scans were complemented by a T2-weighted anatomical scan obtained at the same scanning session. Of these, 248 were successfully pre-processed using the dHCP anatomical pipeline, generating brain tissue segmentations. In addition, brain masks (released with the dCHP anatomical data) and cortical quasi-probabilistic segmentations (Data and code availability, Resource 2) of anatomical scans generated using a 3D U-Net based tool (Uus et al., 2023) were available for all cases with an anatomical scan.

### 2.2. Overview of data processing stages and data structure

Figure 1 graphically presents the overview and inter-dependencies of processing stages, derived data, and complementary data/images utilised in the process. The processing stages comprise of MB-SENSE image reconstruction, dynamic distortion correction, motion correction, and temporal filtering. The naming convention for the outputs of each stage, along with pointers to their location within the data structure, is available on the online resource (Data and code availability, Resource 3).

**Figure 1.**
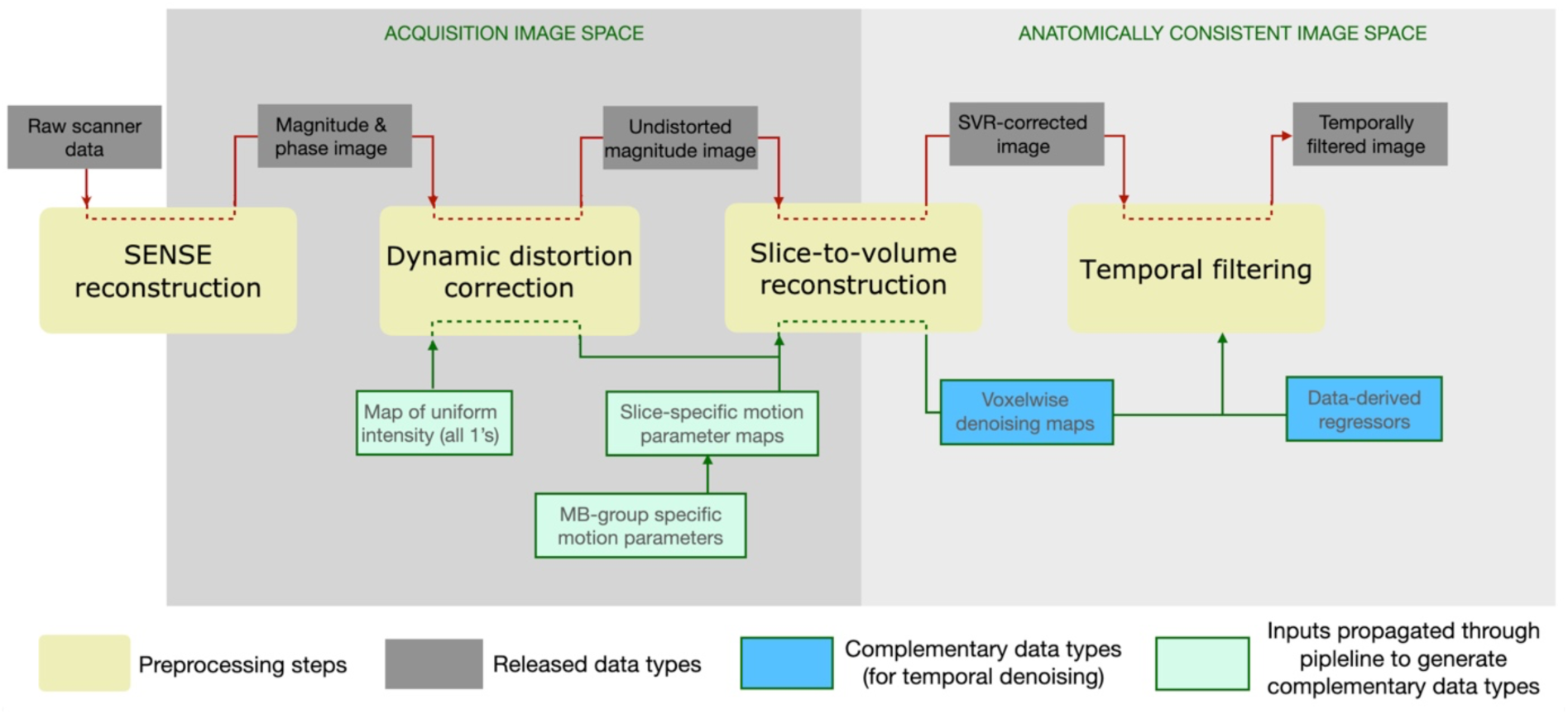
Overview of the processing stages and data structure of fetal dHCP fMRI.

Of relevance, the released data contains items in two imaging spaces: the native acquisition space and the anatomically consistent (tissue) space. The anatomically consistent space results from distortion and motion correction processes, which involve warping, moving, and rotating the original image slices, so that the resulting voxel timeseries represent the temporal signal evolution at specific tissue locations.

### 2.3. MB-SENSE image reconstruction

We used the soft SENSE reconstruction proposed as part of ESPIRIT (Uecker et al., 2014) for considering motion or fat-shift induced model inconsistencies, which was extended to account for MB acquisition (Zhu et al., 2016). Sensitivities were obtained from a single-band (SB) dataset with matched readout (included in the release). Nyquist ghosting correction parameters were obtained for all slices in the field-of-view by using the calibration information collected with the SB data. Three image components were reconstructed with soft SENSE, one corresponding to the target reconstruction and the other two to artefactual information. Spatially adaptive regularisation maps (Fuderer et al., 2004) were constructed for each image component by combining SB reconstructions and the corresponding eigenvalue maps from ESPIRIT.

### 2.4. Spatial Corrections

#### 2.4.1. Dynamic distortion correction

A major challenge of fetal imaging is overcoming the effects of fetal motion and its interaction with the changing *in utero* and maternal environment. The presence of gas bubbles in the gut, changes in maternal body pose during respiration and other incidental movements all cause susceptibility-induced B0 inhomogeneities resulting in highly unpredictable spatial and temporal signal fluctuations, particularly in the fetal brain boundaries. Therefore, spatially and temporally resolved dynamic shot-by-shot B0 field correction is required for improved imaging, especially when imaging at 3T where the aforementioned inhomogeneities are amplified. As a result, the efficiency of registration-based methods (Hutter et al., 2018; Kuklisova-Murgasova et al., 2018; Oubel et al., 2012) may be compromised in this scenario.

We instead opted to use the phase information in the reconstructed gradient echo EPI images acquired for fMRI, as it allows direct estimation of dynamic distortion and therefore separation of distortion and motion correction problems. As residual B0 dynamic evolution is proportional to the phase of the observed signal, its estimation can be posed as a phase unwrapping problem. The solution is obtained by a global phase unwrapping method based on a 4D weighted least squares formulation (Ghiglia & Romero, 1994) with weights constructed by the combination of magnitude information and local phase gradients. A global method is used due to its robustness to local deviations of the phase-based distortion model due to structural noise. Its application has been effective in removing clear distortions in many individuals, with an example shown in Figure 2A.

**Figure 2.**
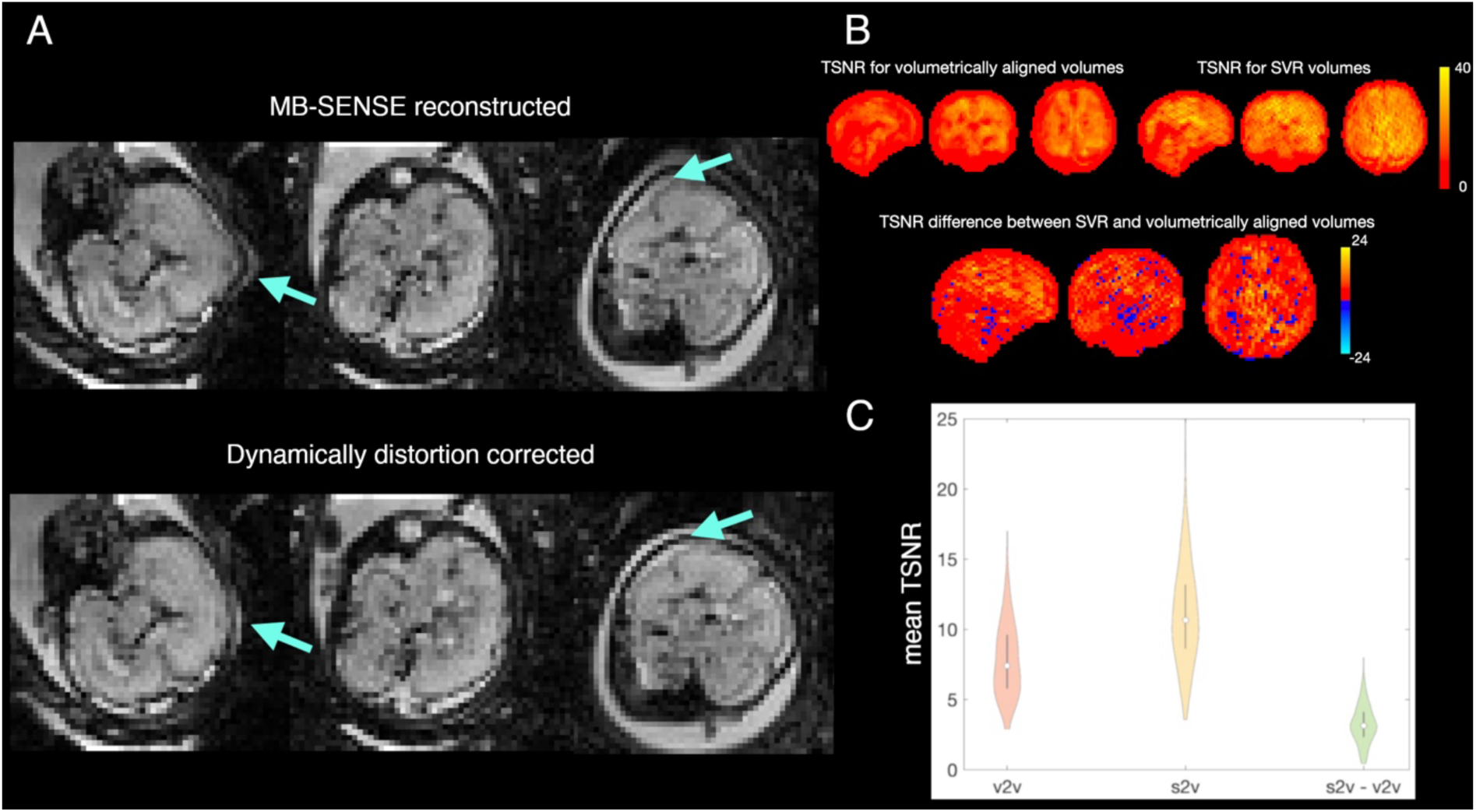
Spatial corrections in the fetal dHCP dataset. Images are in radiological orientation (right is left). A) Distortion correction: Visual example of image data before and after dynamic distortion corrections; B) Motion correction: TSNR for volumetrically aligned, slice-to-volume reconstructed (SVR) volumes and the difference between the two for an exemplar case. Positive values for the latter indicate higher TSNR for the slice-to-volume approach; C) Motion correction: Distributions of averaged (across image) TSNR in the entire dHCP fMRI dataset; v2v – volumetric alignment, s2v – slice-to-volume approach, s2v-v2v – the difference between slice-to-volume and volume-to-volume approaches (s2v-v2v). Positive values for the latter indicate higher TSNR for the slice-to-volume approach.

#### 2.4.2. Motion correction

FMRI data can be severely compromised by motion between the acquisition of different slices. Estimation of rigid-body fetal head motion at the slice level is challenging and correction is ill-posed in the absence of orthogonal stacks (as are typically acquired for anatomical imaging). To address these challenges, we have followed a multi-scale strategy. First, volume-to-volume motion estimates are obtained by standard multiresolution longitudinal registration. Then, these are used to initialise slice-to-volume motion estimation and correction with motion states defined jointly for simultaneously excited slices. Motion parameters are obtained by registering the collected information corresponding to each motion state against a motion-compensated temporal average target. In a final step, motion compensated reconstructions of each fMRI volume are obtained by using the conjugate gradient algorithm for inverting a forward model of the observed fMRI data that considers previously estimated motion parameters and second order regularisation in the slice direction. The whole procedure is based on the framework introduced in (Cordero-Grande et al., 2018b).

The advantage of using slice-to-volume alignment compared to volume-to-volume alignment (still the default approach in much ex utero image pre-processing) can be observed both at individual and group levels. Figure 2B shows the map of temporal signal-to-noise ratio (TSNR, calculated as signal mean divided by its standard deviation over timeseries) difference between both approaches for an exemplary individual. Figure 2C shows average-across-map group-level TSNR distributions for each approach and their difference, indicating the benefit of the slice-to-volume alignment.

### 2.5. Native-to-template mappings

At this stage, the spatially corrected fMRI data were integrated into the dHCP volumetric registration infrastructure (Figure 3), adopting an approach previously implemented for the neonatal dHCP data release (Fitzgibbon et al., 2020). This allows users to flexibly manipulate the data and map between template space and native spaces of all modalities. The remainder of this section will focus on the detailed description of how this infrastructure has been built. A reader interested in processing functional data only can proceed to Section 2.6, without missing out on important details.

**Figure 3.**
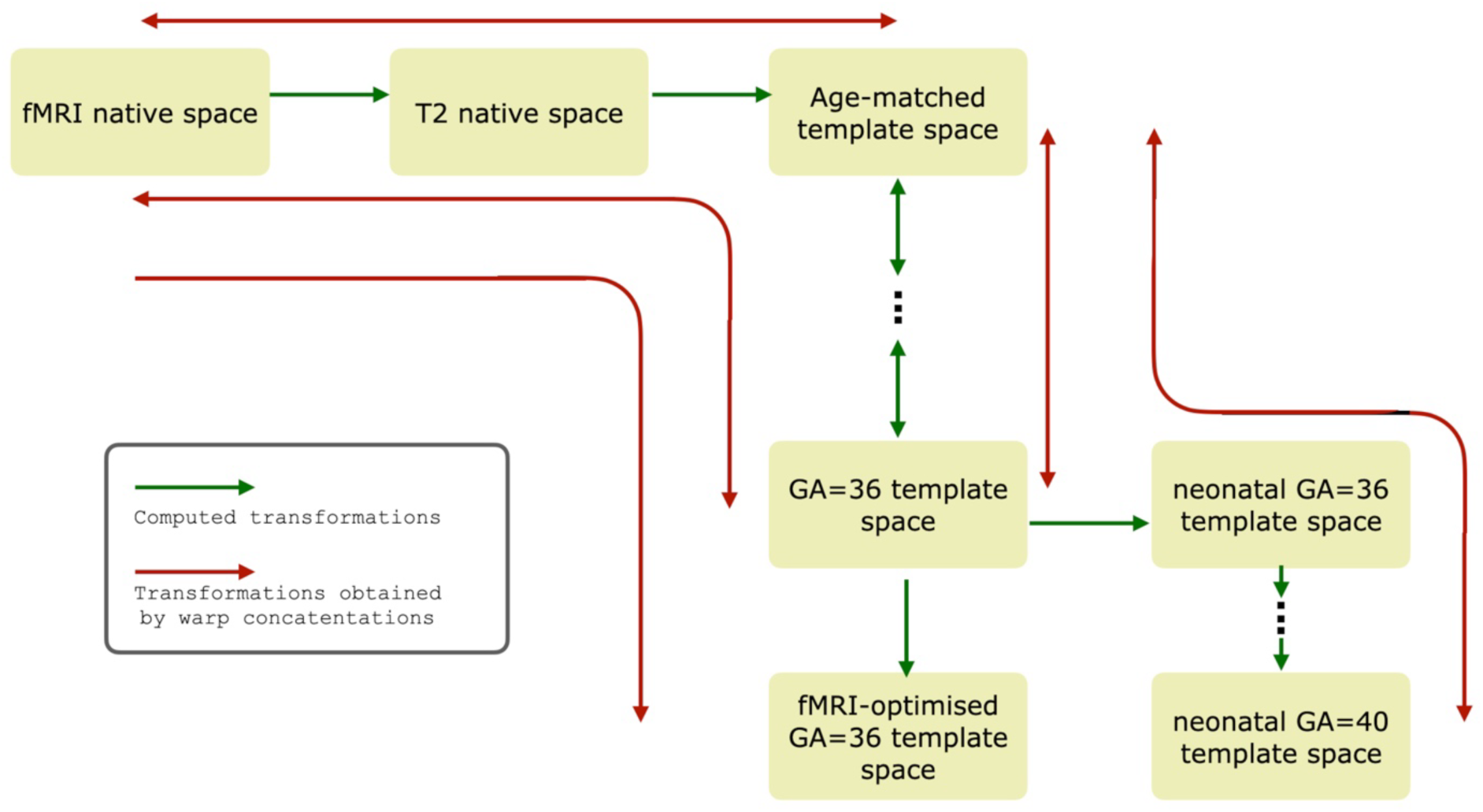
Registration infrastructure associated with the fetal dHCP fMRI data. Concatenated transformations between native fMRI and template spaces are part of the release. Concatenated transformations between template spaces (within-fetal and fetal-to-neonatal) are available at g-node (Data and code availability, Resource 2).

The building blocks of this infrastructure are: 1) linear mapping between native (i.e., individual) functional and T2 spaces, 2) non-linear mapping between native T2 and age-matched template spaces, and 3) non-linear mapping between each pair of age-adjacent templates. The first two building blocks are included in the release, whereas the between-template mappings are available at the dHCP fetal weekly structural atlas repository (Uus et al., 2023 ; Data and code availability, 2). The age-matched template corresponded to that of the subject’s age when rounded to the nearest integer, except for subjects whose GA was > 36.5 weeks, which were all assigned to the 36-week-old template, the oldest template in the dHCP structural atlas.

All linear mappings, including those that preceded estimation of non-linear transformations, were computed using FSL FLIRT. All non-linear warps were estimated using ANTs. The warps were converted into the FSL *fnirt* format, which can be concatenated using the *convertwarp* tool from the FSL library (Smith et al., 2004) in order to create composite displacement warps, incorporating both linear and non-linear components. These composite warps would provide mappings between distant imaging spaces while avoiding multiple image interpolation steps.

To lower the computational burden for the dataset users, two types of composite warps and their inverse have been made available in the release: 1) mapping between native functional and age-matched template spaces and 2) between native functional and 36-week-old template spaces, which could be used for the group-level data synthesis. In addition, the release includes a forward mapping between native functional and the 36-week-old template space optimised for the group-level fMRI analysis, the creation of which is described Section 2.5.4. Details of the procedures used to obtain the required mappings are included below.

#### 2.5.1. Native functional to structural mapping

For cross-modal mapping between native fMRI and T2 spaces, a modified standard procedure, as implemented in the FSL’s *epi_reg* tool, was used. It consists of two stages: a) rigid whole-brain linear registration (using global search and mutual information as a cost function), followed by b) refinement using boundary-based registration (BBR) with local absolute differences cost function and restricting the search angle to ±20°. All other parameters were set to default.

As the fetal brain is surrounded by maternal tissues, including high-intensity amniotic fluid which has an intensity similar to that of cerebrospinal fluid (CSF), accurate masking is required to ensure good alignment, particularly for whole-brain registration. The loose masks used for motion correction of fMRI data, obtained using a spherical Markov Random Field deformable model based on (Cordero-Grande et al., 2011), were employed to initialise alignment.

Subsequently, iterations were performed between function-to-structure alignment and back-projection of high-quality T2 masks into the native functional space.

The details of the procedure are as follows. The alignment of a functional mean image to an anatomical scan was initialised using the masks used for fMRI motion correction. The results were visually checked for being approximately correct and, if necessary, were recomputed after manual mask adjustments. The following steps were then repeated twice. First, anatomical masks were projected into the functional space by inverting the mapping obtained from the initial alignment. These new masks were applied and the N4 corrections for intensity non-uniformity (bias field) within the new mask were applied to the mean fMRI image. The corrected image was then used to re-compute the rigid alignment.

The output from the second iteration of the above procedure was then used to initialise BBR. For the cases (N=26) which lacked the white matter (WM) segmentation required for BBR, binarised cortical quasi-probability maps were used instead, thresholded by minimising the difference between the volume of the thresholded maps and the volume of the cortical mask obtained by the dHCP anatomical pipeline segmentation in the rest of the sample. The results of whole-brain and BBR were visually checked and compared against each other by one of the co-authors (VRK). The BBR alignment was retained if it was rated (based on visual inspection) as performing similarly or better than the whole-brain alignment.

Out of 273 cases with an anatomical scan, 8 scans failed the function-to-structure alignment. All of these failed cases had very poor MB-SENSE reconstruction quality and thus were dropped from the subsequent pre-processing. The BBR alignment was retained in 254 out of 265 cases. The differences between the two approaches were typically subtle but non-negligible in many cases, as in an example of whole-brain and boundary-based registrations presented in Figure 4A.

**Figure 4.**
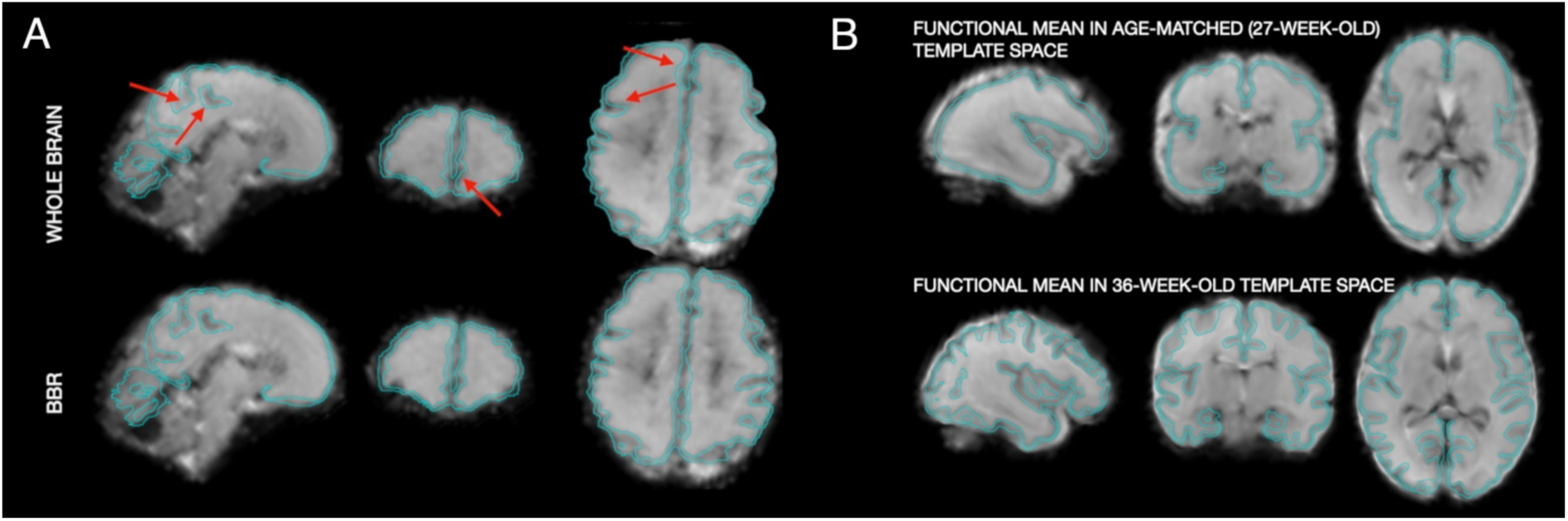
Mapping between image spaces. A. Example of whole-brain vs boundary-based mapping between native fMRI and T2 spaces. Images are in radiological orientation (right is left). The outline of the native structural cortical segmentation is overlaid in cyan. Compare regions pointed by arrows following the whole-brain registration and corresponding regions following BBR. B. Example of mapping the mean functional image into age-matched (27-week-old) and 36-week-old template spaces. The outline of the template cortical segmentation is overlaid in cyan. The original functional data are shown in Figure 2A. Note an accurate alignment of this young case with the “old” (36-week-old) template, despite large morphological differences, owing to concatenation of transformations between age-adjacent template spaces.

#### 2.5.2. T2 to age-matched template mapping

The registrations between native T2 images and age-matched templates were initialised by 12-degrees-of-freedom linear registration using FSL FLIRT (Jenkinson et al., 2002). Default parameters (including cross-correlation metric as a cost function) were utilised. Non-linear alignment was obtained using ANTs SyN diffeomorphic multi-channel registration (Avants et al., 2011), with T2-weighted and cortical probability maps serving as moving sources (i.e., the images to be aligned) and corresponding maps of the dHCP fetal template as the targets.

Local cross-correlation metric was used as a cost-function. Default parameters, as implemented using ANTs *antsRegistrationSyN.sh* wrapper function, were utilised.

The motivation for using separate tools for linear and non-linear parts of registration was that FLIRT demonstrated a robust performance for functional-to-T2 alignment across all ages whereas ANTs is considered to be a state-of-the-art tool for estimation of non-linear mapping (Klein et al., 2009) and proved to perform reliably in the neonatal dHCP cohort (Fitzgibbon et al., 2020). All outputs were inspected visually, and we found no cases of apparent misregistration. The script performing registration between native T2 and age-matched template spaces is available at Resource 3 (Data and code availability).

#### 2.5.3. Template-to-template mapping

The registration procedure between age-adjacent fetal templates (and between fetal and neonatal 36-week-old templates described below) mirrored the registration procedure for individual T2 images to the age-matched template, i.e., was initialised with FLIRT and non-linearly refined with ANTs using T2 and cortical probability maps as registration channels. The older template in each pair was used as the target. The example of mapping a functional image into age-matched and 36-week-old template spaces is shown in Figure 4B.

#### 2.5.4. Fetal-to-neonatal template mapping

To create the mappings between dHCP neonatal (Serag et al., 2012; Data and code availability, Resource 4) and fetal templates, a non-linear transformation was computed between fetal and neonatal 36-week-old templates, followed by creation of composite warps to achieve a direct mapping between arbitrary neonatal and fetal GA template spaces. For the purpose of this study, the mappings between each week fetal template and the “standard” 40-week-old neonatal template were explicitly computed (Data and code availability, Resource 2).

#### 2.5.5. fMRI-optimised standard space for group-level analyses

To compensate for possible residual distortions or misalignment, the registration of the fMRI data to the standard group space was further optimised for group-level analyses. For this sake, mean-across-time native fMRI data were mapped into the 36-week-old template space and a grand-average (i.e., across all subjects) of the mapped data was computed, as well as the grand averages of T2 images and WM and cortical segmentations. We then used ANTs to run a multi-channel non-linear alignment. The subject’s fMRI mean and the grand-average of all subjects’ fMRI means were used respectively, as moving sources and target for the first channel of alignment and the cross-subject means of T2 volumes, and WM and cortical segmentations were used as both moving sources and targets in three other channels. In other words, the deformations were driven by the fMRI image intensities, with identical structural anatomical features between the moving source and the target used to provide spatially varying (contingent on the anatomy) constraints for the scale of deformations allowed. The weight for each structural data channel was assigned to be 10% of the fMRI data channel. The obtained warps were concatenated with warps mapping functional data to the 36-week-old template, allowing for a direct mapping between native fMRI and standard fMRI-optimised spaces.

### 2.6. Temporal filtering

As the standards for temporal filtering of fetal fMRI are yet to be established, the development of a regression-based temporal filtering model was accompanied by an in-depth study of the effect that various types of deconfounding regressors may have on data characteristics, thereby promoting a better understanding of signal properties. We started with an identification of sources for prominent artefacts in the fetal fMRI data and split them into 3 coherent classes. We then selected candidate deconfounding regressors for each class and defined an order in which they were incorporated into a temporal filtering model. After temporal filtering, the data underwent quality assessment, identifying cases suitable for group-level analysis. Finally, we performed a post-hoc study of the unique effects that each set of deconfounding regressors had on the data metrics. For the latter, we used TSNR and seed-to-brain correlations, with the left and right thalami as seeds.

We define 3 broad classes of artefacts. The first class pertains to factors that may affect spatial similarity of the volumes across time. Broadly speaking, here we deal with the effect of imperfections in spatial corrections on the evolution of the signal in the temporal domain.

Potential artefacts attributed to this class are: a) poor motion correction in the presence of large fetal motion; b) signal leakages not fully suppressed during MB-SENSE reconstruction; c) residuals of dynamic distortion correction; d). low-frequency temporal drifts in signal intensity, related to gradient system instabilities.

The second class of artefacts constitute motion-induced changes in the temporal evolution of the signal, potentially causing biology-unrelated covariance structure. Within this group, particular attention is required for spin history artefacts. Their manifestation is likely to take the form of spatially non-stationary image intensity changes (*travelling waves*) (Ferrazzi et al., 2014), which are particularly difficult to address using traditional spatial ICA denoising (which presumes spatial invariance of the signal source).

The third class pertains to the sequential MB data sampling scheme used in the dHCP which induces variable temporal gaps between the acquisition of two spatially adjacent slices. In all cases except for the first and last slices from two adjacent MB stacks, this gap corresponds to the time in between two consecutive RF excitations. In contrast, the temporal gap is on the order of the repetition time (TR) for the slices at the edges of the MB stacks, which can result in abrupt signal changes between them.

These broad classes of potential artefacts lead to the usage of three groups of temporal deconfounding regressors, specific to each class.

#### 2.6.1. Group 1. Temporal regressors for spatial inconsistency artefacts

This group consisted of detrending regressors, volume censoring (“scrubbing”) regressors (Power et al., 2012) and two novel types of voxelwise regressors, which we call folding and density maps respectively. Together they constitute a minimal set of denoising regressors which were used both in combination with other regressors to define more complex denoising models and on their own to residualise the data prior to obtaining data-derived regressors using spatial ICA (see below, Group 2 regressors).

*Detrending regressors.* These were formed by the first 10 columns of the discrete cosine transform matrix, which for the current data corresponded to a high-pass filter with cut-off at 0.0067Hz (150 secs).

*Volume censoring (“scrubbing”) regressors.* This was implemented by spike regressors (Lemieux et al., 2007; Satterthwaite et al., 2013), which in effect equates a censored volume to a temporal average. The criterion for censoring was based on the spatial similarity of a volume to that average. As a metric for the latter, we utilised the root mean square difference (RMSD) measured in terms of the ratio to the grand average of data intensity. More precisely,

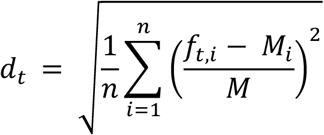

where ***f_t,i_*** is the image intensity in voxel location ***i*** , ***i*** = 1, 2, . . , ***n*** , and volume ***t*** , ***M**_i_* is the temporal median of ***f_t,i_***, and ***M*** is the spatial mean of ***M_i_***.

A volume *t* was censored if ***d_t_*** − ***median***(***d_t_***) > .05. The threshold was determined empirically in order to capture not only global perturbations in image characteristics due to excessive motion but also identifying the volumes with strong local artefacts, caused either by incomplete volume sampling or strong leakage artefacts. The volumes that immediately followed an above-threshold volume were also censored, due to higher likelihood of spin-history impacting the integrity and motion parameter estimates of the next volume (Friston et al., 1996).

*Folding maps*. We utilised the timecourses of the voxels that were simultaneously acquired with a target voxel as deconfounding regressors for potential leakage artefacts in that voxel. For this, we applied a positional shift to the fMRI timecourses, such that the timecourse of a voxel simultaneously acquired with the timecourse of a target voxel was assigned to the latter voxel’s location. Given MB = 3, two 4D maps of this type were formed, which we called folding maps.

As noted previously, there is a distinction between acquisition and anatomically consistent spaces (as in Figure 1). For this reason, distortion and motion correction operators must be applied to the folding maps in order to bring them from the acquisition space into the anatomically consistent space. As a result, voxels in the resultant folding maps represent weighted averages of the original maps in accordance with the transformations and interpolation schema applied to the data.

*Density maps.* These were designed to remove temporal dependencies potentially induced by the distortion correction procedure. Specifically, in addition to spatial coordinate transformations, the distortion correction involves a Jacobian modulation which compensates for the compression/spread of the signal in the phase encoding direction. Modulation is a voxel-wise multiplicative factor which corrects image intensities and should not, in principle, be correlated with the temporal evolution of the signal after correction; if this is the case it can be assumed that it is likely to be artefactual. To generate the denoising map that carries this information, we created a 4D image of uniform intensity with geometry matched to the geometry of the data in the acquisition space. We then applied distortion- and motion-correction transformations to simultaneously bring the image into the anatomically consistent space and scale it with the applied modulation.

#### 2.6.2. Group 2. Temporal regressors for motion-induced signal changes

This group included two types of regressors: 1) time-courses of non-grey-matter (GM) tissues (WM and CSF), derived in a data-driven manner and 2) 4D voxel-wise maps representing the evolution of motion parameters over time.

*Data-derived regressors.* Data-derived non-GM maps regressors (e.g., Behzadi et al., 2007) were obtained by performing probabilistic ICA (FSL MELODIC) analysis (Beckmann & Smith, 2004) on the data within regions combining subarachnoid and ventricular CSF and WM that were residualised with respect to Group 1 regressors. The mask combining these tissues was eroded by one voxel to reduce partial voluming. For cases in which the dHCP anatomical data failed the quality control assessment, missing segmented WM and CSF masks were generated by projecting the segmentations of age-matched subjects into the native functional space of that particular case and averaging them to produce an approximate map. Depending on the particular denoising model (as described below), the ICA dimensionality was set to 6, 24, or automatically determined using FSL MELODIC implementation of the Minka algorithm (Minka, 2000)/capped at 90 components (whichever was smaller).

*Motion parameter (MP)-based voxelwise maps.* Following slice-to-volume motion correction, each of the 16 MB excitations (i.e., 48 slices / MB factor of 3) in a volume obtains its own set of 6 motion re-alignment parameters. The utility of this information on the motion evolution is compromised by the fact that they describe transformations applied to the slices of the volumes residing in the acquisition space. In other words, each parameter is specific here for a set of 3 slices, but not voxel-specific. To account for this, we created 4D maps for each parameter, where all voxels within slices belonging to a particular MB group were assigned the MP that was used to map them into the anatomically consistent space. After that, the motion-correction operations were applied to these maps to bring them into the anatomically consistent space. The resultant denoising map no longer contains identical values within a slice, since the value for each voxel is derived from interpolation.

12 maps of this type were created. The first 6 corresponded to 3 rotation angles and 3 translation values and the other 6 represented corresponding differentials in the slice direction. The slice differentials were calculated as a difference ***S_t_***_+1_– ***S_t_***, obtained by rearranging the MP timeseries in the anatomically consistent space, ***V***_1_, ***V***_2_, … , ***V***_350,_(where any ***V_k_*** is a ***n*** × 1 vector, ***n*** is the number of voxels, and 350 is the number of timepoints), into the simultaneously excited slices-to-slices timeseries ***S***_1_, ***S***_2_, … , ***S***_350, × 16_ (where any ***S_k_*** is a ***n***/16 × 1 vector, with 16 the number of excitations in the acquisition of a volume, i.e., they comprise slices in the same MB excitation). The reason for creating maps of simultaneously excited slices’ differentials was to enrich MP-based regressors with dynamics in the intrinsic dimension for spin-history effects.

The above maps were used to create a broader set of MP-based regressors, unique for each voxel. By defining the temporal evolution of the set of 6 MP and their differentials in the slice direction at a voxel in location ***i*** as 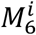 and 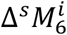, respectively, and first- and second-order differentials in volume-to-volume direction as Δ^1^ and 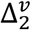, respectively, the complete set of 60 regressors for each voxel was defined as 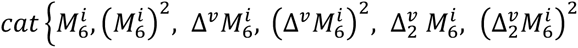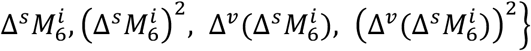, where ***cat***{} denotes column-wise concatenation. This set was transformed first by z-scoring each column and then by using principal component analysis, considering only non-censored volumes, with the number of components retained to account for 99% of variance in the regressor set.

*Selection of Group 2 regressors – signal implanting test.* Having a broad choice of Group 2 regressors potentially creates an overfitting problem, especially when considering that the temporal filtering model can combine both MP-based and data-derived regressors. Overly aggressive temporal filtering can inadvertently remove the biologically relevant signal fluctuations of interest, which could seriously hamper further analysis given the inherently low SNR of fetal fMRI data, especially at the single-subject level.

To guide the selection of the best performing denoising models, we designed a surrogate test, which we call the signal implanting test. This test is based on injecting a biologically-plausible signal into the data, and observe which composition of temporal regressors allows for: 1) most accurate recovery of this signal and 2) maximally preventing signal loss. For this, we utilised the neonatal group-level maps (Fitzgibbon et al., 2020) and their timecourses, estimated by regressing the group maps against individual data in the neonatal subjects from the dHCP neonatal cohort, aged between 37 and 43 weeks. The timecourses were paired at random with the fetal subjects and the group maps were projected into the fetal native functional spaces. The product of the timecourses and the group maps was then scaled to either 3% or 6% of the temporal standard deviation of the real data for each particular subject and added to the data, constituting an “implanted signal”. The data were then residualised with respect to a candidate set of denoising regressors, and the spatial maps were regressed against the denoised data to obtain estimates of their timecourses; the correlation between their vectorised product and vectorised implanted signal were taken as a single measure of the quality of signal recovery. Similarly, the same procedure was applied to the timecourses that were removed from the data with implanted signal during denoising. A high correlation between removed and implanted signal would indicate that the model inadvertently removes signal from the data. We will refer to the latter measure as signal loss statistics.

We tested several models that differed in the number of data-derived and MP-based regressors (Figure 5A). All candidate models were complemented with Group 1 regressors. Figure 5B shows the results of the test. Overall, the analysis indicates that MP-based denoising models with a limited number of non-GM data-derived regressors performed best with respect to the signal recovery statistics. Considering both signal recovery and signal loss statistics, the MP-based model with 6 non-GM regressors (highlighted) performed the best. Firstly, it performed better than the others with respect to signal recovery statistics. Secondly, the model also showed a similar performance to the model with 0 non-GM regressors with respect to signal loss statistics whereas the model with 24 non-GM regressors showed significant losses.

**Figure 5.**
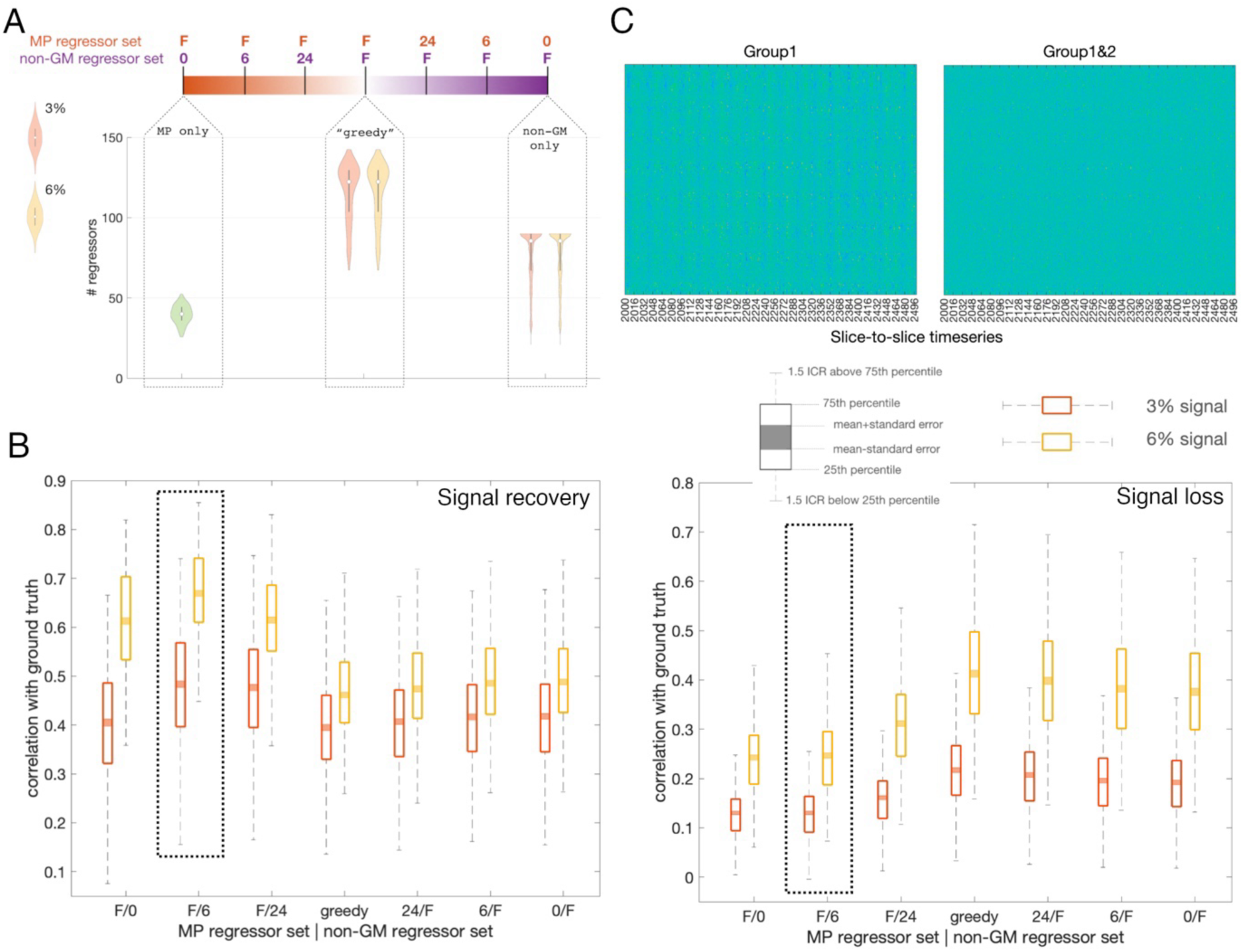
Group 2 regressors. A) Scope of temporal denoising models assessed using the signal implanting test. “F” stands for the “full” set for a particular (MP-based or data-derived) type of regressors. Correspondingly, the “F/0” model includes full set of MP-based and no data-derived regressors, the “0/F” model includes the full set of data-derived and no MP-based regressors, with intermediate models in between. The “greedy” model includes maximal number of regressors of both the MP-based and data-derived type. The sample distributions for the number of regressors in the extreme and greedy (F/F) models are plotted underneath, calculated for 266 scan sessions that passed the function-to-structure alignment. As the full set of data-derived regressors is estimated from the data, implanting different volumes of signal (3% or 6%) may change the output of the Minka algorithm/capping at 90. The sample distribution of MP-based regressors is calculated from the spatial means, given that .99 variance may be represented by a different number of components across voxels. B) Results of the signal implanting test. Model ordering as in A. Left - signal recovery statistics. Right - same for the signal loss statistics. The best performing model is highlighted. C) Segment of slices-to-slices timeseries before (Group1 regressors only) and after application of Group 2 regressors. Ticks along x-axis mark TRs, i.e., 16 slices. Note the vertical stripes, repeating at the rate of approximately 2-3 TRs and likely to be caused by mother’s breathing cycle, that are significantly reduced after application of Group 2 regressors.

In three cases, the application of the best-performing model to the real data resulted in a numerical overflow. These cases were of a very poor quality and had a large number (> 210) of volumes to censor; for this reason, no attempt was made to correct the issue retrospectively. Consequently, the fully processed dHCP dataset consists of 263 sessions (247 unique individual subjects, 132 male, 113 female, 2 unrecorded).

Visual inspection of the filtered data reveals a notable reduction of the travelling waves pattern, indicating that MP-based voxel-wise regressors are effective in ameliorating the spin-history effects. Quantification of this phenomenon is not straightforward, but it can be observed qualitatively when temporally demeaned data are rearranged into slices-to-slices timeseries (Figure 5C).

#### 2.6.3. Group 3. Temporal regressors for sampling scheme artefacts

A set of regressors addressing artefacts related to the difference in temporal separation between first and last slice of adjacent MB stacks compared to other neighbouring slices were derived using single-subject spatial ICA with dimensionality set to 30 components run on the timeseries following their residualisation with respect to the model that included Group 1 and (best performing) Group 2 regressor sets. Prior to this, the data were slightly smoothed using a FWHM=1 mm 3D Gaussian filter. Here we sought to identify slab-like spatial patterns repeating roughly across the MB stacks width (i.e., 16 slices). Two observers (VRK and DB) independently performed the rating of spatial component maps, with three scores allowed: ‘to remove’, or ‘to be equivocal’, or ‘to keep’. One observer started the rating from the first subject in the subject list, and the other started the rating from the middle subject. To ensure that similar subjective criteria were applied across the cohort, each observer used the rating of the first 20 subjects to get accustomed to the task. The ratings for these subjects were reviewed again after the observer completed the rating of the whole dataset. The criterion for consensus to regress out component timecourses was defined as either ‘to remove’ by both raters or ‘to remove’ by one and ‘to be equivocal’ by the other.

Figure 6A shows examples of components that were rated by consensus as representing a sampling artefact. The average number of such components per subject was 7.6, sd = 3.96 (Figure 6B). Residualisation of the data with respect to the timecourses of these components was performed on top of residualisation with respect to Group 1 and Group 2 regressors.

**Figure 6.**
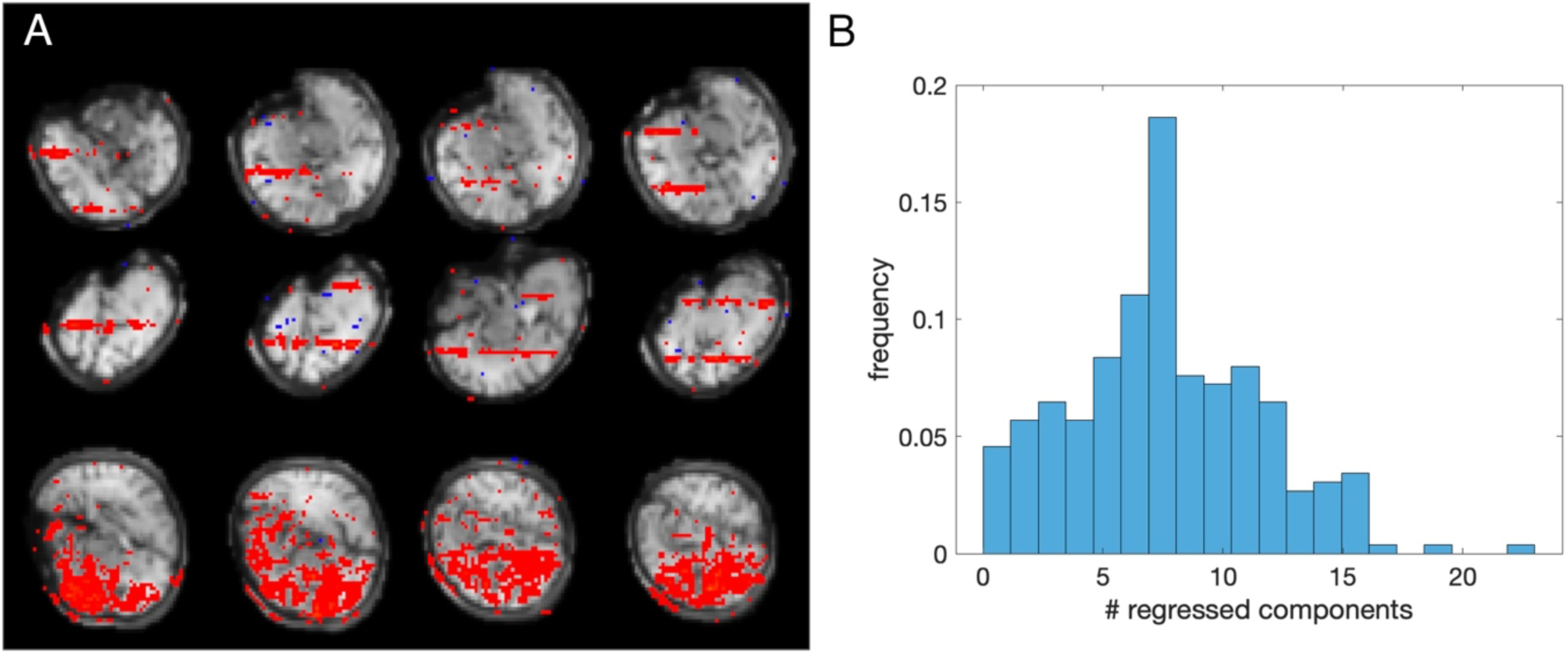
Group 3 regressors. Images are in radiological orientation (right is left). A) Examples of components with double- slab-like spatial structure in an exemplary subject; B) Distribution of the number of components rated to be removed per subject

Because the timecourses of the independent components are not constrained to be orthogonal, the regression coefficients for the timecourses to regress out were estimated in combination with other components’ timecourses. This procedure generated the most thoroughly processed data in the release.

#### 2.6.4. Quality control and data selection for group-level analyses

To rate the quality of individual pre-processed data and select cases suitable for group-level analyses, we used both visual assessment and parametric measures.

For visual assessment, the timeseries were rated by an observer (VRK) using a 5-point scale, 0 – fail, 1 – bad quality (seemingly unusable), 2 – seemingly usable, 3 – reasonably good, 4 – good. This was complemented with the two parametric measures, mean (over voxels) *tsnr* and *dvars*, disregarding censored volumes. Specifically, mean-over-voxels *tsnr* was defined as:

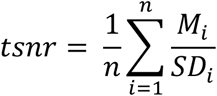

where:

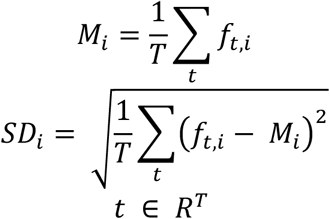

and

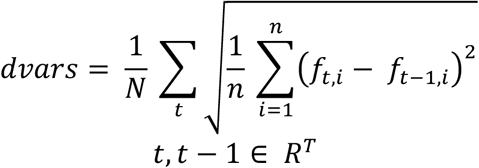

where ***t*** is the order number of a volume, ***R***^2^is the set of ***T*** uncensored volumes, ***N*** is the number of consecutive pairs of uncensored volumes, ***t***, ***t*** − 1 ∈ ***R***^2^, and ***n*** is the number of voxels. Both measures were calculated in the functional native space, within a binarised anatomical segmentation image, which provides a tight mask by excluding non-brain tissues (e.g., subarachnoid CSF).

The decision rule was as follows: all cases rated >1 (high score) were considered as suitable for analysis EXCEPT when they were EITHER positive *dvars* outliers OR negative *tsnr* outliers with respect to all cases rated > 1. Specifically, the outlier ***O**_dvars_* for *dvars* and ***O**_tsnr_* for *tsnr* were defined as:

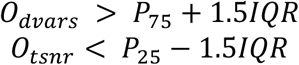

where ***P***_25+_ and ***P***_75_ are 25^th^ and 75^th^ percentiles of the cases rated > 1, and ***IQR*** is their interquartile range.

Furthermore, all cases rated <=1 (low score) were also considered as suitable for an analysis, if BOTH their *dvars* AND *tsnr* were, respectively, < ***P***_75_ AND > ***P***_25_ of cases rated > 1. All ratings as well as their *dvars* and *tsnr* measures are available at Data and code availability, Resource 3.

Figure 7A-B shows that visual assessment of the pre-processed data, *dvars* and *tsnr* were all in reasonable agreement. Based on the predetermined criteria, 217 out of 263 scans passed the QC (114 male, 101 female, 2 unrecorded). Figure 7C demonstrates an example of the QC report, available for each individual subject (Data and code availability, Resource 3).

**Figure 7.**
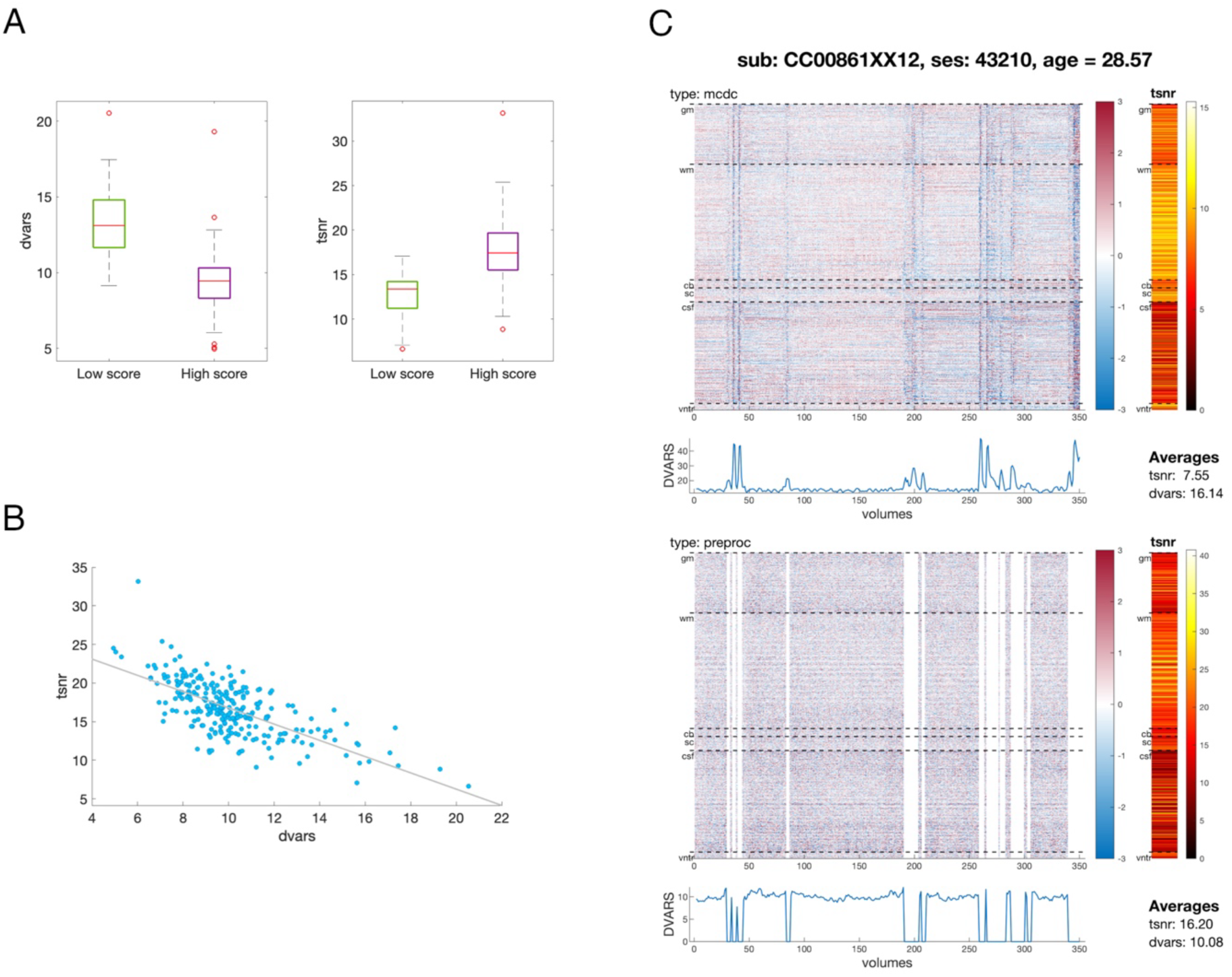
Quality assessment. A) Average dvars and tsnr for high (> 1) and low (<=1) scores in visual quality assessment. B) Relationship between average tsnr and dvars; C) QC report for an exemplar subject, showing carpet plots of z-scored intensity, spatial profiles of TSNR and temporal evolution of dvars for the data after motion and distortion correction (type: mcdc, without censoring) and for the fully processed data (type: preproc, including censoring).

#### 2.6.5. Group-level statistics of deconfounding regressor groups

Figure 8 shows the summary statistics of all 263 fully pre-processed cases for the temporal filtering regressor groups with respect to the loss of the effective degrees-of-freedom (DOFs, = number of regressors, Figure 8A) and TSNR (Figure 8B). On average, the fully pre-processed data lose approximately 32% of effective DOFs, and gain more than 2 times TSNR, compared to the data that underwent spatial distortion and slice-to-volume motion correction only. The loss in DOFs is significantly higher than reported in adult imaging (e.g., approx. 15% for minimally pre-processed data in the Human Connectome Project (HCP) - Smith et al., 2013), but much of it occurs via volume censoring, which has no analogy in HCP.

**Figure 8.**
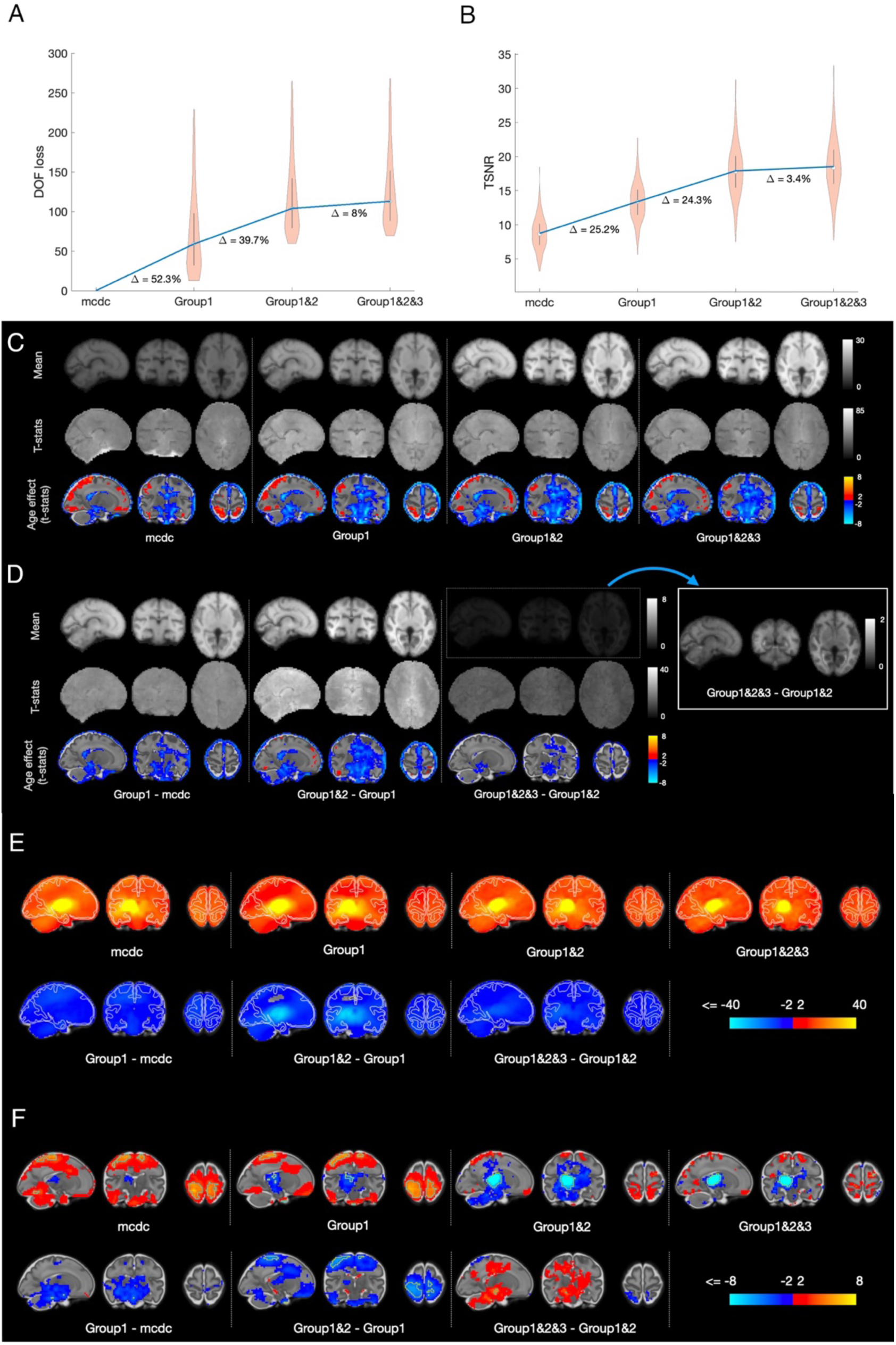
Statistics for different temporal filtering models. Images are in radiological orientation (right is left). A) The loss in effective DOFs. “mcdc” signifies a null model, i.e., motion and distortion corrected data, without temporal filtering. The average per model is represented by its median. Differences in percentages are calculated with respect to the average for Group1&2&3 model. B) The gain in average TSNR. The average per model is represented by its mean. C) Spatial maps of TSNR and estimates of the age effect; Right is left. D) Spatial maps of differences in TSNR between different denoising models. E) T-statistics for the intercept of seed-to-brain thalamic correlation maps (top row) and the maps representing the difference in correlations after applying different denoising models (bottom row); hemisphere on the left is uni-lateral to the location of the seed; F) same as E) for the age effect.

In order to evaluate the local effects of the temporal filtering models, we utilise the dataset capacity for group-level analyses, afforded by the released registration infrastructure, projecting TSNR maps (smoothed with a FWHM = 3 mm Gaussian kernel in the native space) into the fMRI-optimised group space. The aligned maps were then modelled with linear regression, with demeaned age as a co-variate, providing estimates of average TSNR characteristics in the sample and their association with age.

The mean TSNR maps (Figure 8C) and the maps of the TSNR differences (Figure 8D) between temporal filtering models help appreciate the effect of each regressor group. The patterns are presented in terms of fitted beta coefficients for intercept and after conversion into t-statistics for both intercept and age effect. In terms of beta coefficients, both the average TSNR and its gain from application of the more complex denoising model was the highest in WM, which reflects the fact that mean intensity (i.e., numerator in the TSNR formula) is higher in the WM than in the GM and CSF. This points to the multiplicative effect of noise, supported by the observation that WM TSNR is no longer greater than GM TSNR when converted into t-statistic values, taking into account across-subject variability.

As TSNR is not informative with respect to the effects of deconfounding regressors on estimates of brain functional connectivity, we investigated the latter in the context of seed-to-brain correlation maps, using the left and right thalami as seeds. The analysis was performed in the sample of individuals GA > 24.5, which passed quality control criteria. This comprised 201 subjects (106 male, 93 female, 2 unrecorded). Individual correlation maps were computed for the left and right thalamus separately in the native functional space (following FWHM = 3 mm smoothing) and projected into left-right symmetrised space (Data and code availability, Resource 3) for group-level analysis. To enable computing of the left-right subject’s average maps, the maps of the left thalamus connectivity were left-right flipped. We then modelled the average maps using linear regression with demeaned age as a covariate.

Figure 8E-F shows the results for the fitting. In summary, using a more complex denoising model results in a decrease of correlations throughout the brain. However, perhaps with exception for Group 1 regressors (spatial correlation −0.20 between “mcdc” and “Group1-mcdc” maps in Figure 8E), this decrease was proportionally scaled, i.e., more prominent in areas showing a stronger correlation to the seed timecourse prior to model application (spatial correlation −0.95 between “Group1” and “Group1&2-Group1” maps and −0.73 between “Group1&2” and “Group1&2&3-Group1&2” maps). A similar proportional effect was observed for the age effect (Figure 8F; spatial correlations = −0.04, −0.92 and −0.62, respectively). The most profound decrease for both intercept and age-effect was observed when including Group 2 regressors.

### 2.7. Group-ICA analysis

To demonstrate the dataset capacity for multi-variate network analysis, we performed estimation of functional modes (“networks”) in the same sample as in the analysis of thalamic connectivity. All individual data were smoothed in native space using a FWHM=3 mm Gaussian kernel. To account for heterogeneity inherent to a sample with large maturational differences, we first concatenated the timeseries of age-matched individuals (as defined for age-matched template registration) and compressed them to 1400 principal components for each age. After transforming within-component (across voxels) values by z-scoring, the components were concatenated with similarly defined components from other age groups and compressed to a sample-average set of 1400 principal components which was then fed into FSL melodic ICA to obtain whole-sample decomposition into 25 components. In other words, the procedure ensured a balanced contribution of each age to the final factorisation, whereas z-scoring of values within each component ensures that factorisation attends to common patterns irrespective of their absolute scale, thereby implicitly taking into account that these patterns may have different prominence across ages.

Figure 9A shows functional modes obtained in these data, which predominantly comprise of components localised to one brain region (hereafter - dominant node). Despite this, there was a notable prevalence of component pairs (7 pairs, grouped together in the figure), where dominant nodes are a left-right reflection of each other. Out of these 7 pairs, 3 pairs (ICs 14 vs 21, 10 vs 16, and 6 vs 19) showed symmetry not only in the location of a dominant node, but also in the location of secondary non-negligible nodes. Another notable observation is that 3 components (ICs 15, 20, and 24, highlighted in bold green in Figure 9A) were characterised by a node pair, one dominant and another smaller, located in the homologous regions of the two hemispheres, and another (IC 3) showed one medial component with approximately equal bilateral representation.

**Figure 9.**
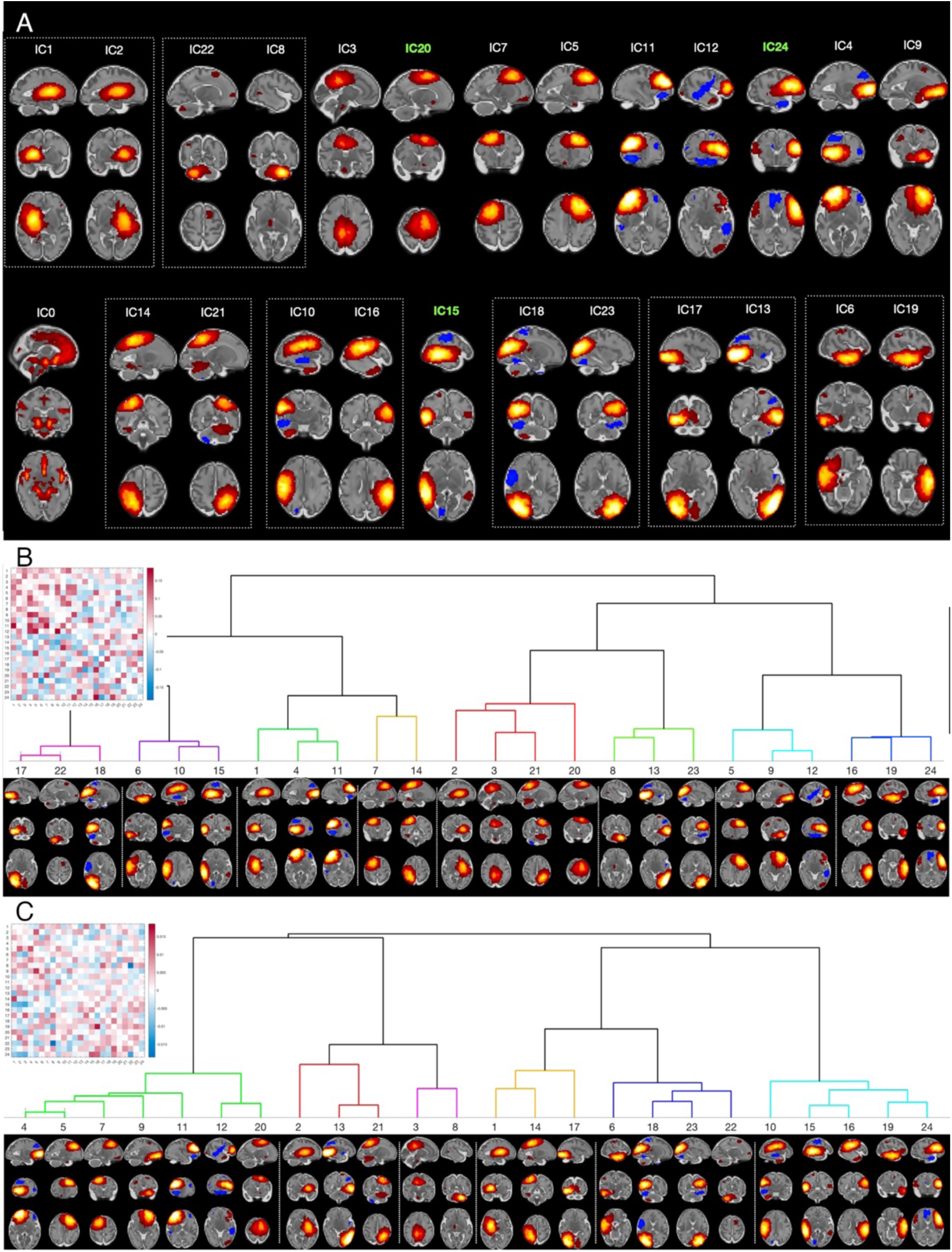
Modes of covariation (fetal “resting-state networks”). Images are in radiological orientation (right is left). A) Spatial maps grouped by their anatomical location. The dotted-line boxes show pairs of maps showing a distinctive interhemispheric symmetry. Components with non-negligible bilateral homotopic representation are highlighted in bold green. B) Full (lower triangle)-partial (upper triangle) correlation matrix and hierarchical clustering based on full correlations transformed into t-statistics. C) Same as B) for the age-related changes in correlations.

To evaluate the structure of whole-brain functional architecture, we derived the temporal time courses of the obtained 25 ICs by spatial regression (Beckmann et al., 2009) against individual fMRI data in native functional space. For each subject individually, we computed the correlation matrix between component timecourses, excluding timecourses of IC0 which is of a distinctively vascular origin. We then combined matrices from all subjects in order to fit each element with linear regression, using demeaned age as a covariate. The procedure resulted in two matrices, one containing estimates of an average strength of association between each pair of components’ timecourses and the other providing estimates of their age-related changes. The elements of these two matrices were converted into t-values, thereby accounting for between-subject variability. Normalised Laplacian embedding of each matrix (positively thresholded to make this estimable) was then used to represent the relationships between components in terms of a Euclidean distance between their coordinates in the embedded space. We then applied hierarchical clustering using the Ward method (Ward, 1963), based on the coordinates in the first 3 non-null dimensions.

The results of clustering (Figure 9B) demonstrate a weakness of association between contralateral nodes in terms of average strength of their correlation; the overall tendency for this metric was to form clusters of unilateral nodes, sometimes representing a symmetrical image of each other (e.g., ICs 17,22,18 vs ICs 8,13,23 and ICs 6,10 vs ICs 16,19) and/or representing coherent functional systems, such as right dorsal fronto-parietal (ICs 7,14), leftdominated sensori-motor (ICs 2,3,20,21), left (ICs 5,9,12) and right (ICs 1,4,11) frontal.

Clusters that mix both uni- and contralateral components are more evident in the analysis of age-related changes (Figure 9C). Of particular interest is a cluster (ICs 10, 15, 16, 19, 24) that combines areas belonging to semantic and language processing. Notably, it includes only a left but not right anterior temporal node, which aligns well with a previous report on connectivity of these areas in the adult brain (Hurley et al., 2015).

## 3. Discussion

Despite its rapid progress (De Asis-Cruz et al., 2021; Ferrazzi et al., 2014; Jakab et al., 2014; Ji et al., 2022; Karolis et al., 2023; Schöpf et al., 2012, 2014; Seshamani et al., 2016; Sobotka et al., 2022; Taymourtash et al., 2021, 2023; Thomason et al., 2015; van den Heuvel et al., 2018), fetal fMRI remains a relatively novel field, where pre-processing standards are yet to be established and which until now lacked an open-access resource available to researchers. In this paper, we introduce an open-access fetal fMRI data repository and present a systematic approach for pre-processing, spanning stages from image reconstruction to group-level analysis. We implemented state-of-the-art methods for the spatial reconstruction and correction of the data, including MB-SENSE reconstruction, dynamic distortion and slice-to-volume corrections, and a tailored temporal filtering model with attention to the prominent sources of structured noise. The final dataset consists of 263 fully processed cases, with 217 cases determined to be suitable for the connectivity analyses. We thereby provide both a resource and a high-quality template to facilitate further development and benchmarking of methods for fetal fMRI. In the remainder of the discussion, we focus on the potential areas for improvement.

Studying fetal samples using fMRI is perhaps an exemplary case where the age of a subject simultaneously represents the most important variable-of-interest and the most prominent confound. The latter manifestations encompass, to name a few, signal-altering differences resulting from rapidly changing tissue properties, drastically different morphology, and three-fold differences in effective resolution. These factors entail various challenges for the progress of the fetal fMRI field. Firstly, they create obstacles for reproducible experiments.

The standard practice of scanning one participant repeatedly (Duff et al., 2022; Noble et al., 2017; Shah et al., 2016; Shehzad et al., 2009) in fetal imaging settings is practically complicated and ethically dubious, as this approach can render meaningful results only if there is no significant temporal gap between scanning sessions. Secondly, they entail difficulties in defining sound quantitative metrics, appropriate for the entire age range, that would enable comparative testing between various processing approaches. Thirdly, they may entail differences in magnitudes of distortion and motion, thereby affecting data quality in an age-related manner. The released data, that includes outputs from different processing stages and information on motion and dynamic field mapping, represents therefore a valuable asset for investigating motion patterns at different gestation ages and dynamic distortion and motion relationship. Finally, they also complicate the inter-subject data fusion for group-level analysis and defining ground truths that could guide pre-processing approaches in order to harmonise data quality across ages and fit a broad range of potential goals for a study. As an example, the immature fetal WM is thought to contain its own functional units within temporary developmental structures such as the subplate and constitutes a metabolically active area, especially at younger ages (Colonnese & Phillips, 2018; Tolner et al., 2012). Consequently, studying the WM BOLD signal at younger ages may require a different approach to the temporal filtering problem.

Our approach to the temporal filtering follows the trend that was set by the seminal work by (Power et al., 2012) and (Satterthwaite et al., 2013), evaluating the effect of motion on resting-state connectivity in adults. This work demonstrated an improved precision of connectivity estimates following application of sophisticated denoising models enriched with motion-related or data-derived deconfounding regressors (Power et al., 2014; Pruim et al., 2015; Satterthwaite et al., 2013; Yan et al., 2013). Here, by virtue of using the recovery of an artificially implanted cortical signal as a metric to assess efficiency of temporal filtering models, we sought to optimise the data for the analysis of cortical networks. We were conscious of the possibility that the usage of complex models may inadvertently remove relevant signal information, i.e., that selectivity may be prioritised at the expense of sensitivity, which may be suboptimal in the fetal fMRI low-SNR setting. Namely, we found that the data-derived regressors (obtained using ICA on non-GM timeseries), which represent a popular choice for *ex utero* data (Kiviniemi et al., 2003; Kochiyama et al., 2005; Salimi-Khorshidi et al., 2014) including paediatric data (Fitzgibbon et al., 2020), are particularly prone to undesired implanted signal removal in the fetal fMRI.

Correspondingly, our approach to temporal filtering relied on a rich set of MP-based regressors, helping us to achieve a notable reduction of artefacts, including difficult-to-tackle spin-history artefacts. In spite of taking these measures, we observed proportionally scaled decreases in TSNR and estimates of thalamic connectivity following the application of complex MP-enriched models. These appear to suggest that in real data signal and noise do not fully comply to the additivity assumption of the signal implanting test. The proportional-like decrease in the strength of the age effect following application of MP-based and data-derived regressors points to the same possibility. A case in point here is the age-related changes in connectivity between the thalamus and sensorimotor and parietal areas. On the one hand, age-related increases in these functional connections are expected based on current knowledge of prenatal brain development from post-mortem studies (Kostović et al., 2019; Kostović & Judaš, 2010). On the other hand, these changes are best detectable in the data before any temporal filtering, indicating contribution of age-related structured noise to this contrast. Taken together, these findings suggest several far-reaching implications. Firstly, they suggest that magnitude of the structured noise may scale with age and therefore the strength of the age effect cannot be accepted as a sole criterion for optimisation of the model for temporal filtering. Secondly, in the absence of a ground truth, the intricate entanglement of signal and noise complicates the determination of the temporal filtering complexity for a right sensitivity-selectivity trade-off. Thirdly, more sophisticated motion modelling could be required to increase selectivity of temporal regressors to structured noise.

We therefore invite dataset users to develop temporal filtering models that could better fit the requirements of their analyses and/or investigate the ways to boost their statistical power, including alternatives not considered in this paper, such as more aggressive spatial smoothing and lowpass temporal filtering. To facilitate this, we release a rich set of complementary data in the form of denoising maps and provide Python code that can be used with the regressors in the released database or adapted to include new ones (Data availability statement, Resource 3). We also remark that the implementation of data reconstruction and spatial corrections was developed to be compatible with proprietary format of raw data collected by the scanner used in the dHCP, considered not only fetal fMRI but many other MRI scanning domains, and was optimized for the local computing hardware infrastructure. Therefore, we have decided not to provide a ready-to-use code release for data reconstruction and spatial corrections to be run externally but we release the code for dynamic correction (Data and code availability, Resource 5).

Another area for improvement relates to incorporation of deep learning models for improving brain extraction, spatial corrections and temporal denoising in the proposed pipeline, with recent approaches mature enough, in particular, for the brain extraction problem (e.g., (Rutherford et al., 2022)). A particularly challenging scenario is encountered in scans where the brain’s orientation varies significantly in time. In the current implementation, such differences could result in an orientation flip in some volumes of the reconstructed timeseries, which had to be censored for downstream processing. The extreme situation where a fetus may turn upside-down within an fMRI scan is far from being manageable with current analysis techniques, which exemplifies that data rejection criteria and quality control also merit future efforts for improvement and standardisation across studies.

Finally, detailed investigations are also needed to determine how the large age-dependent variability in brain characteristics can analytically be incorporated into building group-level normative models of functional brain development. Fetal fMRI data reveals idiosyncratic properties (Karolis et al., 2023), apparent in various forms, including when results of group-level ICA factorisation between fetal and neonatal samples are contrasted. Analysis in the fetal brain typically reveals a dominance of a single-node functional modes and a paucity of spatially distributed patterns (Ji et al., 2022; Karolis et al., 2023), whereas in the neonatal brain the latter are not uncommon, especially those left-right symmetrically organised (Eyre et al., 2021; Fitzgibbon et al., 2020; Gao et al., 2015). This may reflect a genuine property of the fetal brain’s functional organisation, its immaturity and perhaps the functional state of the brain while *in utero*, but may also simply be a reflection of a lower signal-to-noise ratio or may suggest that the method, that implicates estimation of a group-level ‘mean’ model in a developmentally heterogeneous sample, is not appropriate for these data (Karolis et al., 2023). In this paper we adopted pre-processing steps in group-ICA analysis that aimed to ameliorate the challenges of data fusion across ages. The obtained results were more prone to revealing signatures of distributed spatial patterns than previously reported (e.g., Ji et al., 2022; Karolis et al., 2023). Furthermore, they reveal a range of symmetrically located network pairs, tentatively pointing to an emergence of a bilaterally organised functional architecture (Thomason et al., 2013). However, despite these elements of qualitative convergence with the neonatal data, harmonisation of in- and ex-utero cohorts for a combined analysis remains a task for future research.

In conclusion, research into functional brain development in utero is strongly motivated by a large body of evidence that shows the fetal period has critical importance for individual development. This information can be obtained non-invasively using fetal fMRI, but much work is required in order to establish the standards of data acquisition, pre-processing and analysis in this challenging domain. The current dataset represents a unique resource that aims to provide a firm foundation for advancing fetal fMRI from its current status as a niche research field to its deserved and timely place at the forefront of the community-wide efforts to build a life-long connectome of the human brain.

## Data and code availability

The data and code presented in this paper are available in the following locations:

1. dHCP fetal release: https://nda.nih.gov/edit_collection.html?id=3955
2. The dHCP volumetric template can be accessed at: https://gin.g-node.org/kcl_cdb/fetal_brain_mri_atlas. Folder “structural” contains T2-weighted spatiotemporal atlas, folder “composite_warps” contains fsl-style warps between different ages of template; folder “composite_warps_fetal2neonatal” contains mappings between fetal template and the 2 neonatal templates spaces. Folder “cnn_cortex_probability” contains quasi-probability maps of cortical segmentations.
3. Materials associated with this paper, such as sample demographics, naming convention, supplementary data, QC reports, the script to perform registration of native T2 images to age-adjacent templates, and the python code used for performing temporal filtering, are available at: https://gin.g-node.org/kcl_cdb/dHCP_fetal_fMRI_release_paper.
4. The neonatal templates and mappings between different ages are available at: https://git.fmrib.ox.ac.uk/seanf/dhcp-resources
5. The code for distortion correction is available at: https://github.com/mriphysics/fetalPhaseEPI/releases/tag/1.0.0. Enquiries about sharing the existing code for image reconstruction and motion corrections and guidance on its application should be directed to co-author LCG.

## Ethics

The study was approved by the UK National Research Ethics Authority (14/LO/1169). Written informed consent was obtained from all participating families prior to imaging.

## Conflict of interests

The authors declare no conflict of interests.

## Author contributions

Conceptualisation – V.R.K., L.C.G., A.D.E., T.A., S.M.S, E.D., J.H.

Writing - Original Draft - V.R.K., L.C.G. Investigation - E.H., A.P., M.A.R.

Software – V.R.K., L.C.G., A.U., D.P., M.D., A.M.

Data curation – V.R.K., L.C.G., V.K., N.H., D.P., D.B.

Methodology – V.R.K., L.C.G., S.P.F., A.U., J.W.M., S.W., D.C., M.P., M.D., L.Z.J.W., E.C.R., A.M., S.R.F., J.O.M.

Formal analysis – V.R.K., L.C.G., V.K., A.U. Visualisation – V.R.K.

Validation – V.K.R., L.C.G.

Funding acquisition - D.R., A.D.E., T.A., S.M.S., J.H. Resources – A.D.E., S.M.S., J.H.

Project administration – T.A., J.H. Supervision – T.A., S.M.S, E.D., J.H.

Writing - Review & Editing – V.R.K., L.C.G., J.W.M., S.W., D.C., M.D., L.Z.J.W., E.C.R., S.R.F., T.A., E.D., J.H.

## Acknowledgments

The Developing Human Connectome Project was funded by the European Research Council under the European Union Seventh Framework Programme (FP/20072013)/ERC Grant Agreement no. 319456. The Wellcome centre for Integrative Neuroimaging is supported by core funding from the Wellcome Trust [203139/Z/16/Z]. The authors also acknowledge support in part from the Wellcome Engineering and Physical Sciences Research Council (EPSRC) Centre for Medical Engineering at King’s College London [WT 203148/Z/16/Z], the Medical Research Council (MRC) Centre for Neurodevelopmental Disorders [MR/N026063/1], and the Department of Health through an NIHR Comprehensive Biomedical Research Centre Award (to Guy’s and St. Thomas’ National Health Service (NHS) Foundation Trust in partnership with King’s College London and King’s College Hospital NHS Foundation Trust). VRK and TA were supported by a MRC translation support award [MR/V036874/1]. TA was also supported by an MRC Clinician Scientist Fellowship [MR/P008712/1] and a MRC Senior Clinical Fellowship [MR/Y009665/1]. JOM is supported by a Sir Henry Dale Fellowship jointly funded by the Wellcome Trust and the Royal Society [206675/Z/17/Z]. LCG acknowledges funding from MCIN/AEI/10.13039/501100011033/FEDER, EU under Project PID2021-129022OA-I00 as well as Universidad Politécnica de Madrid for providing computing resources on Magerit Supercomputer. SRF’s research is supported by the Royal Academy of Engineering under the Research Fellowship programme [RF2122-21-310]. JWM was supported by PhD funding from the UK Medical Research Council [MR/P502108/1].

## Notes

### Competing Interest Statement

The authors have declared no competing interest.

### Summary of Updates

The updated version includes the revisions following recommendations by the paper reviewers

